# An information-theoretic analysis of the tsetse fly (*Glossina morsitans*) antenna reveals trichoid sensilla as the main carriers of odor identity

**DOI:** 10.64898/2026.10.26.761766

**Authors:** Neeraj Soni

**Author notes:** Corresponding author: (NS).

## Abstract

Tsetse flies locate vertebrate hosts largely by smell, and odor-baited devices are central to tsetse and trypanosomiasis control. A published electrophysiological survey of the *Glossina morsitans* antenna defined seven classes of olfactory sensilla, but descriptive clustering cannot reveal how much information these classes carry or which odors they distinguish. We reanalyzed responses of 182 sensilla to 16–17 odorants using Fisher–Rao information geometry, bias-corrected mutual information, Bayesian decoding, and signal detection theory. In the fruit fly *Drosophila melanogaster*, trichoid sensilla are largely dedicated to pheromones. In tsetse, the three trichoid classes carried more information about odor identity than the four basiconic classes (1.92 vs. 1.14 bits, of a possible 4), a difference of 0.78 bits (95% confidence interval 0.44–1.10), and were strongly excited by the host attractant 1-octen-3-ol. Adding trichoid to basiconic sensilla raised decoding accuracy from 32% to 70% (23% to 55% for decoders tested on held-out sensilla) and identified 1-octen-3-ol on 98% of trials (92% held out). The basiconic code was nearly one-dimensional (participation ratio 1.71), organized along an axis of response strength with 1-octen-3-ol at its extreme. The CO_2_-sensitive ab4 class was the most sensitive detector for 11 of 16 odorants, whereas ab2 carried no detectable odor information. Classes shared up to 0.41 bits of their information with one another, an overlap only weakly predicted by the similarity of their mean tuning. On a shared odor panel, *Drosophila* receptors carried twice the information per channel and, with channel number matched, spanned about 1.5-fold more dimensions. These conclusions held in simulations at the real sample sizes, under alternative noise models, and when classes were defined on independent odorants. The tsetse antenna thus combines redundant, detection-oriented basiconic channels with trichoid sensilla that carry host-odor identity, principles relevant to the design of odor baits.

**Author summary:** Tsetse flies spread sleeping sickness in people and a related wasting disease in cattle, and they find the animals they bite largely by smell. Traps and targets baited with odors are among the most effective tools for controlling these flies, so knowing what the fly’s nose can and cannot tell apart could help us design better baits. Several years ago we recorded how the different smell-sensing hairs on the tsetse antenna respond to a panel of odors. Here we returned to those recordings with a different question: not which hairs respond to which odors, but how much information each type of hair actually carries about which odor is present. Using tools from information theory, we found that one type of hair, which in the fruit fly serves mainly to detect the scents of other flies, carries most of the tsetse antenna’s information about the identity of host odors, including the classic tsetse attractant 1-octen-3-ol. The other hair types act mainly as sensitive alarms that signal that a host odor is present. These results suggest which sensory channels odor baits should be designed to engage.

## Introduction

Tsetse flies (*Glossina* spp.) transmit the trypanosomes that cause human African trypanosomiasis (sleeping sickness) and animal African trypanosomiasis (nagana) across sub-Saharan Africa. Tens of millions of people have lived at risk of infection [1], and nagana imposes a major economic burden on livestock production [2]. Human cases have fallen to historically low levels, and the World Health Organization now targets elimination of *gambiense* transmission by 2030 [3,4]. Vector control has contributed substantially to this progress: odor-baited traps and insecticide-treated targets can reduce tsetse populations by more than 90% [5]. The efficacy of these devices depends on how well they exploit the fly’s olfactory system, which guides host location over long distances [6–8].

We previously carried out a systematic electrophysiological survey of the *Glossina morsitans* antenna [9]. Recordings from 182 sensilla defined three trichoid classes (at1–at3) and four basiconic classes (ab1–ab4), revealed that every class with excitatory responses is excited by the host attractant 1-octen-3-ol, and showed that the ab1, ab3, and ab4 response profiles are strongly correlated. These features distinguish the tsetse antenna from that of *Drosophila melanogaster* [10,11], and they were identified with hierarchical clustering and rank correlation. Such descriptive methods, however, cannot answer quantitative questions that matter for both neurobiology and vector control: How much information about odor identity does each sensillum class carry? How much of that information is shared between classes? Which odors can the periphery distinguish and which does it confuse? Is a class that responds weakly, such as ab2, uninformative or simply tuned to untested compounds?

Information theory and information geometry provide a principled framework for these questions. Shannon mutual information quantifies how much a response reduces uncertainty about the stimulus, independently of any particular decoder [12,13], and its extension to pairs of channels measures how much information they share [14]. Information geometry treats a family of response distributions as a Riemannian manifold whose Fisher–Rao metric measures how distinguishable nearby stimuli are [15,16]. These tools have clarified coding principles in the *Drosophila* olfactory system [17–20] and in theoretical treatments of receptor arrays [21,22], but to our knowledge they have not been applied to the peripheral olfactory code of a disease vector. Here we reanalyze the complete *G. morsitans* dataset [9] with four complementary frameworks: Fisher–Rao geometry of the odor manifold, bias-corrected mutual information, Bayesian decoding, and signal detection theory. A primer on each framework, written for biologists, is provided in S1 Text. We then compare tsetse with *Drosophila* on a common odor panel. The analyses reveal a division of labor on the tsetse antenna: basiconic sensilla provide a sensitive but redundant detection signal, whereas trichoid sensilla, which in *Drosophila* are largely devoted to pheromones [23,24], carry most of the information about the identity of host odors.

## Results

The original survey described the tsetse antenna with bar plots of mean responses, hierarchical clustering of sensilla, and rank correlations between classes [9]. It observed that several classes have correlated response profiles, that 1-octen-3-ol excites nearly every class, that ab2 gives no excitatory responses, and that trichoid sensilla respond to many host odorants. Each of these observations concerns mean responses. What a downstream neuron can learn from a response, however, depends on how different the responses to two odors are relative to how much the response to a single odor varies from sensillum to sensillum. The analyses below incorporate this variability and ask, for each of these observations, how much information it represents.

### The tsetse odor code is low-dimensional, with 1-octen-3-ol at its extreme

We first asked how many independent dimensions the basiconic population uses to represent odors. Principal component analysis of the 143 basiconic sensilla (ab1, n = 68; ab2, n = 17; ab3, n = 51; ab4, n = 7) tested with 16 odorants showed that the first component captured 76% of response variance and four components captured 90.5% (Fig 1C). The participation ratio, an effective count of dimensions [25], was 1.71 of a possible 16. Because sensilla were the samples in this analysis, the low dimensionality partly reflects differences in overall responsiveness between sensilla and classes; the geometry of odor space itself is addressed next.

**Fig 1.**
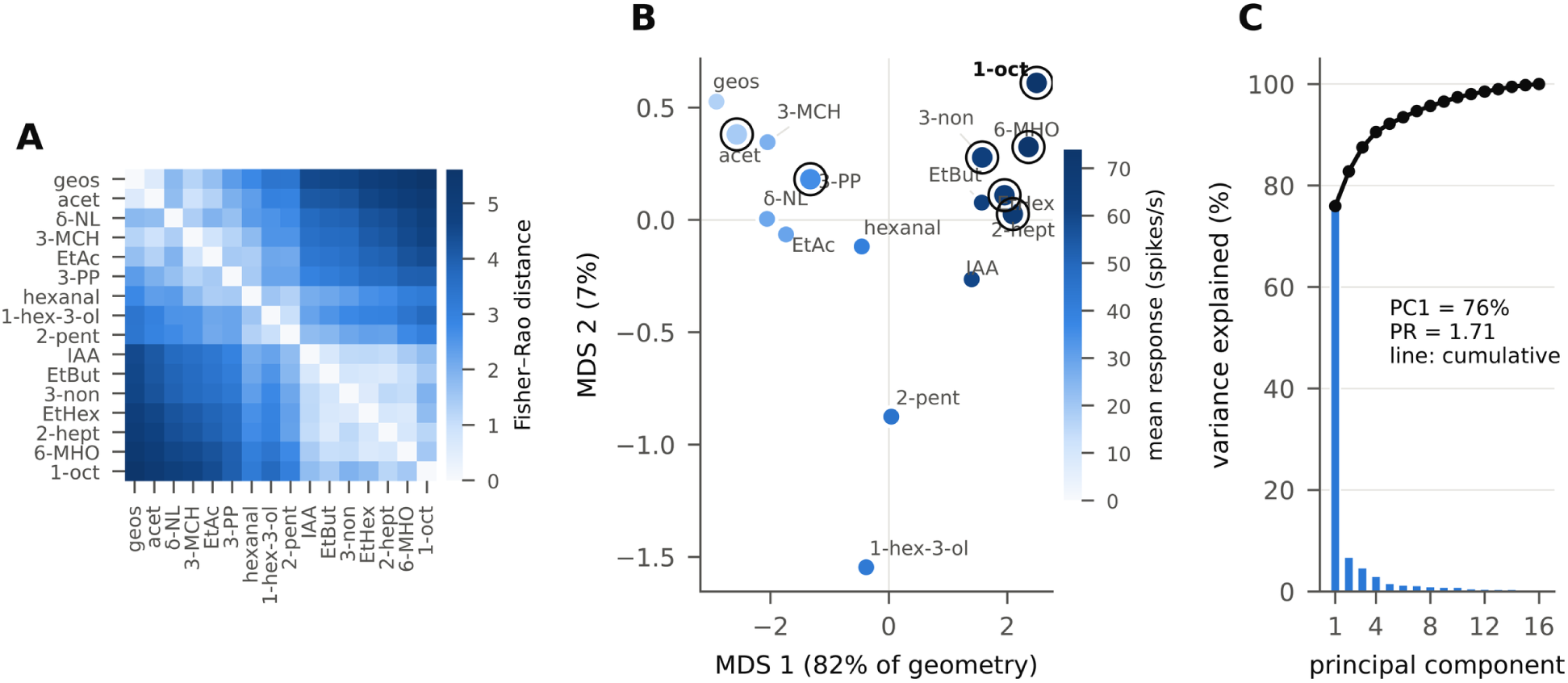
Information geometry of the tsetse basiconic odor code. (A) Fisher–Rao distances between all pairs of 16 odorants, computed on the product manifold of the four basiconic classes (ab1–ab4), each odorant–class response modeled as a Gaussian with the measured mean and standard deviation (SD) across sensilla. Odorants are ordered by their coordinate on the first multidimensional scaling (MDS) axis in B. (B) Classical MDS embedding of the distance matrix in A. Point color indicates the mean response across the four classes; circled points mark odorants associated with vertebrate hosts or used in tsetse baits. MDS axis 1 accounts for 82% of the geometry and correlates with mean response (|r| = 0.997). 1-Octen-3-ol (bold) lies at the strong-response extreme; acetone and geosmin form the closest pair (d = 0.58). (C) Principal component analysis of the 143 × 16 basiconic response matrix (all sensilla, all odorants). Bars, variance explained by each component; line, cumulative variance. PR, participation ratio. Abbreviations: 1-oct, 1-octen-3-ol; acet, acetone; 3-PP, 3-propylphenol; δ-NL, δ-nonalactone; geos, geosmin; 2-pent, 2-pentanol; EtBut, ethyl butyrate; 1-hex-3-ol, 1-hexen-3-ol; EtAc, ethyl acetate; 2-hept, 2-heptanone; EtHex, ethyl hexanoate; 6-MHO, 6-methyl-5-hepten-2-one; IAA, isoamyl acetate; 3-non, 3-nonanone; 3-MCH, 3-methyl-2-cyclohexenone.

To examine discriminability directly, we modeled the response of each class to each odorant as a Gaussian distribution with the measured mean and SD across sensilla, and computed the Fisher–Rao distance between every pair of odorants on the product manifold of the four classes (Fig 1A; Materials and methods). Classical multidimensional scaling of this distance matrix placed 82% of the geometry on a single axis (Fig 1B); negative eigenvalues, which arise because Fisher–Rao distances are not Euclidean, accounted for 0.2% of the total. Position on this axis was almost perfectly predicted by the mean population response to an odorant (|r| = 0.997), so the dominant axis of tsetse odor space was response magnitude rather than odor quality. Because all odorants were delivered at the same dilution, this axis may partly reflect differences in the amount of each odorant reaching the antenna as well as differences in how sensilla respond (Discussion).

The host attractant 1-octen-3-ol occupied the strong-response extreme of this axis. The largest Fisher–Rao distance in the matrix separated 1-octen-3-ol from geosmin (d = 5.60; mean pairwise distance 2.69), and 1-octen-3-ol was the second most peripheral odorant relative to the manifold centroid (2.75 vs. mean 1.96), exceeded only by geosmin at the opposite, weak-response pole. 1-Octen-3-ol was also relatively isolated from its nearest neighbor, 2-heptanone (d = 1.19 vs. a mean nearest-neighbor distance of 0.85). In contrast, the weak-response odorants acetone and geosmin were nearly coincident (d = 0.58, the smallest distance in the matrix). The basiconic code therefore represented odors primarily by how strongly they drive the population, with 1-octen-3-ol as the most distinctive stimulus in the panel.

### Sensillum classes differ markedly in information content

We next quantified how much information each class carried about odor identity. For each class we computed the mutual information I(S;R) between odorant S (16 equally likely odorants) and the response R of a single sensillum, under the same Gaussian model, and subtracted the upward bias expected from finite samples using a shuffle null (Materials and methods) [26]. In simulations from the fitted model at the real sample sizes, this correction removed the upward bias of the plug-in estimate and was, if anything, conservative (Table E in S1 Appendix). Information is measured in bits: one bit is the information needed to answer one yes-or-no question with equally likely answers. Identifying one odor among 16 requires four such questions, so log_2_16 = 4 bits is the ceiling, the information that a perfectly reliable response would provide. A response that carries I bits narrows the 16 candidates, on average, to about 16/2^I^.

Among basiconic classes, ab3 carried the most information (0.80 bits; 95% confidence interval (CI), 0.71–0.88): a single ab3 response narrowed 16 possible odors to about nine. It was followed by ab4 (0.50 bits; 0.05–0.94) and ab1 (0.39 bits; 0.30–0.48) (Fig 2A). ab4 carried less information than ab3 despite producing the largest responses, because its responses varied more from sensillum to sensillum (mean SD 31 vs. 21 spikes/s) and were estimated from only 7 sensilla, hence the wide confidence interval. ab2 carried 0.009 bits (95% CI −0.06 to 0.07), indistinguishable from the shuffle null (P = 0.18, permutation test). Among trichoid classes, at1 (1.09 bits) and at3 (1.05 bits) each carried more information than any basiconic class, and at2 carried 0.30 bits. A nonparametric k-nearest-neighbor estimator that makes no distributional assumption [27,28] gave closely similar values for every class (Fig 2A, diamonds; Table A in S1 Appendix).

**Fig 2.**
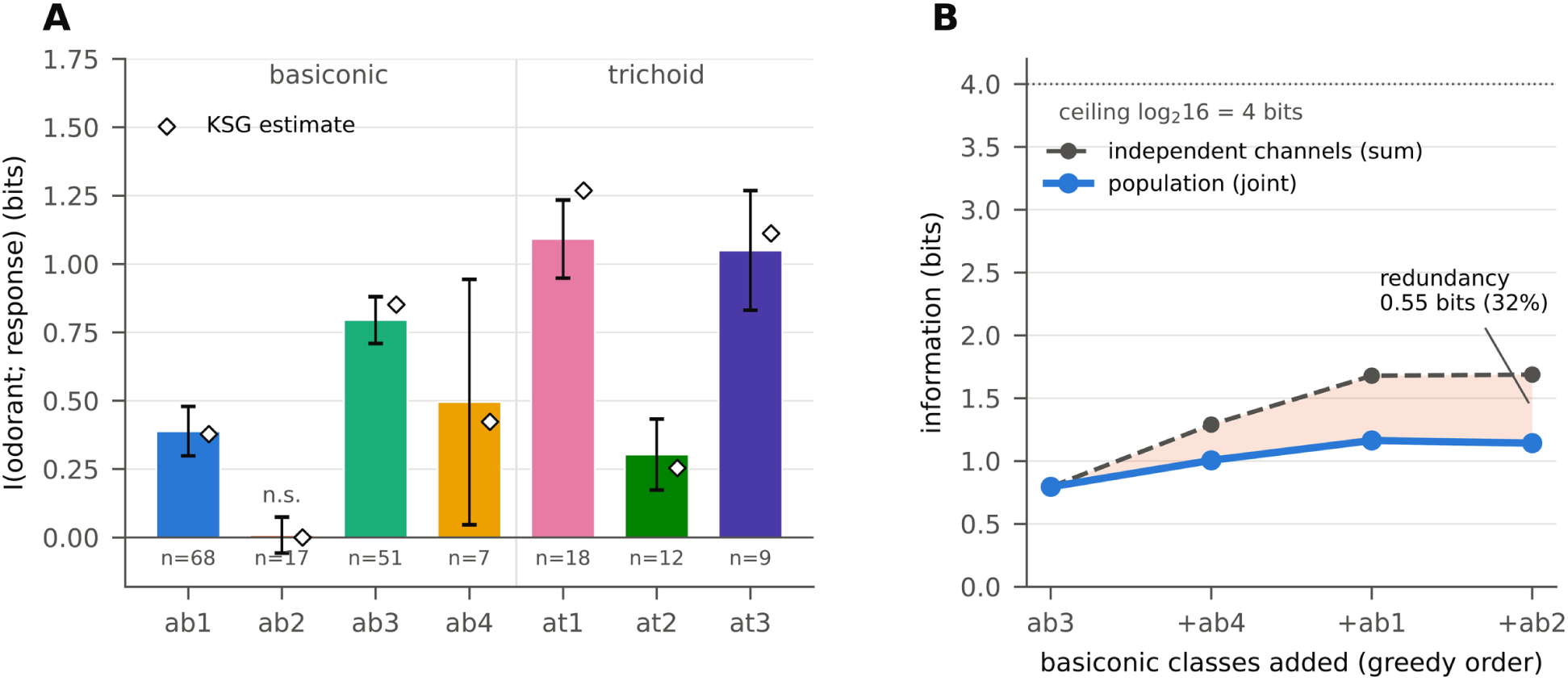
Information carried by tsetse sensillum classes. (A) Bias-corrected mutual information between odorant identity (16 odorants) and the response of a single sensillum of each class. Bars, Gaussian-channel estimate after shuffle correction; error bars, 95% CI from a bias-corrected bootstrap over sensilla; diamonds, nonparametric k-nearest-neighbor (KSG) estimate. n, number of sensilla. n.s., not different from the shuffle null (P = 0.18); all other classes P < 0.01 (100 permutations). (B) Population information as basiconic classes are added in the order that maximizes the gain at each step (solid line) compared with the sum of single-class information (dashed line), which is the value expected if the classes carried independent information. The shaded area is the redundancy. Dotted line, ceiling of log_2_16 = 4 bits, the information needed to identify every odor perfectly.

The information carried jointly by the four basiconic classes was 1.14 bits (95% CI 0.82–1.46), 29% of the 4-bit ceiling; one response from each basiconic class narrowed 16 odors to about seven. Had the four classes carried independent information, their total would have been 1.69 bits. The shortfall of 0.55 bits was redundancy: 32% of the summed single-class information was shared among classes (Fig 2B). Two classes are redundant when, once the response of one is known, the other adds little about which odor is present. In an olfactory system, redundancy lowers the number of odors that can be distinguished but makes the host-odor signal robust: it survives the adaptation or loss of any one class, and it can be averaged across the many sensilla of each class. Adding classes in order of greatest gain showed rapidly diminishing returns: ab3 alone supplied 70% of the population information, ab4 and ab1 together added 0.37 bits, and ab2 added nothing (1.16 bits without ab2, 1.14 bits with it). All seven classes together carried 2.27 bits, 55% of their 4.13-bit summed single-class information; the contribution of the trichoid classes is analyzed below.

### Sensillum classes share much of their information

The original survey noted that ab1, ab3, and ab4, and also at1 and at3, have correlated response profiles, differing mainly in response magnitude [9]. Correlated mean responses suggest, but do not establish, that two classes carry the same information. Whether they do depends on the shape of each profile across odors rather than its height, and on how reliably each class separates the same odors, which in turn depends on response variability; a bar plot of means shows neither. To measure shared information directly, we computed for every pair of classes the interaction information ΔI_ij_ = I(S;R_i_,R_j_) − I(S;R_i_) − I(S;R_j_) [14]. Because the classes were recorded in different sensilla, their joint responses had to be modeled as conditionally independent given the odorant. Under this assumption ΔI_ij_ equals −I(R_i_;R_j_), the information that the two classes’ responses carry about each other through their shared tuning, so it cannot be positive (Materials and methods; we confirmed this identity numerically to within 0.003 bits). ΔI therefore measures redundancy in tuning (signal redundancy); synergy, which would require correlated variability between classes, cannot be assessed without simultaneous recordings. Shared information was substantial (Fig 3A). The largest overlaps linked ab3 with at3 (−0.41 bits), ab3 with ab4 (−0.28), and at1 with at3 (−0.28). The correlated profiles previously noted among ab1, ab3, and ab4 [9] were thus reflected in substantial shared information, and the overlap extended across morphological types: trichoid at3 was more redundant with basiconic ab3 than with any other class. Every pair involving ab2 had |ΔI| ≤ 0.03 bits, consistent with ab2 carrying essentially no odor information to share.

**Fig 3.**
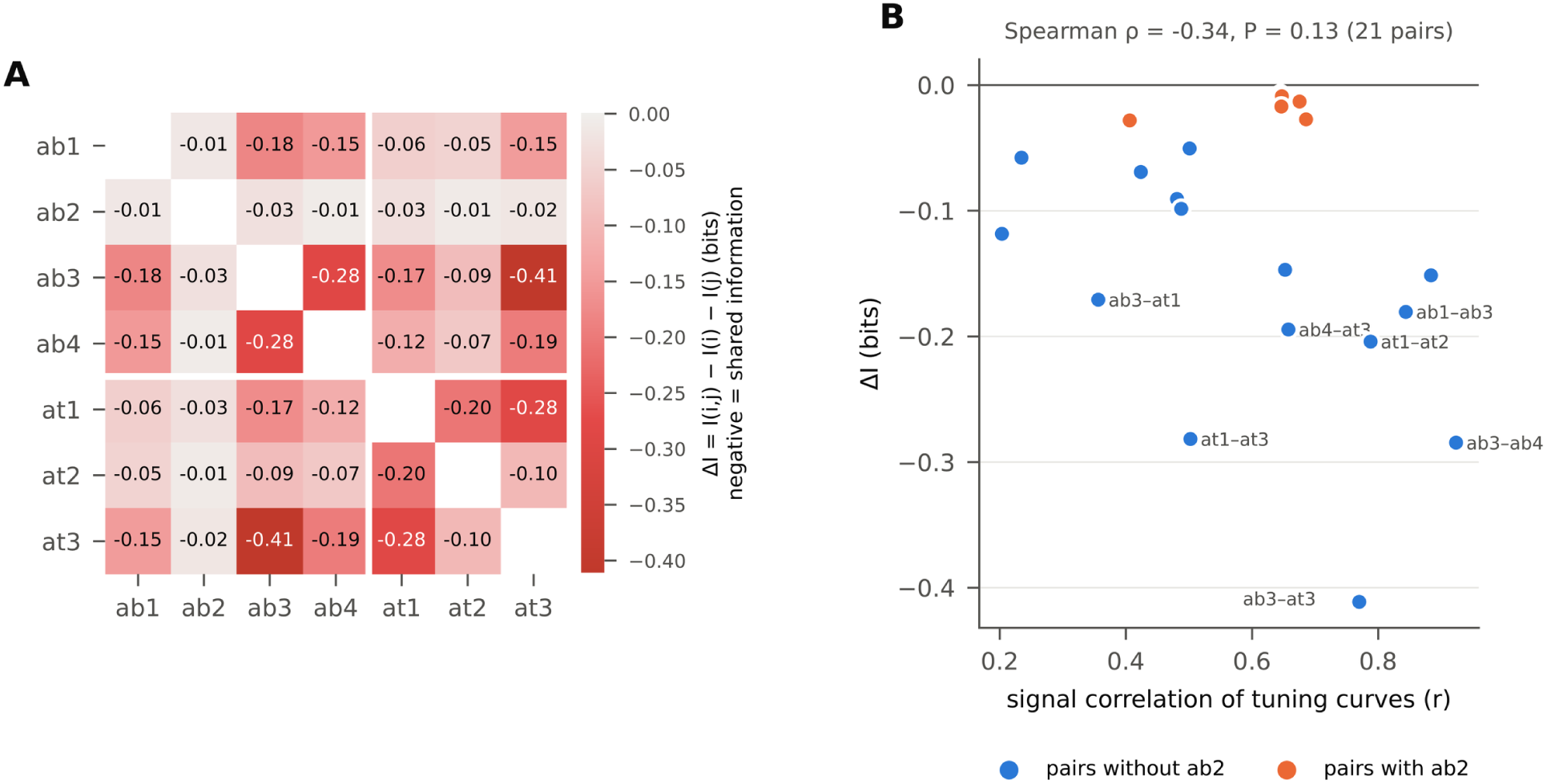
Redundancy among sensillum classes. (A) Interaction information ΔI_ij_ = I(S;R_i_,R_j_) − I(S;R_i_) − I(S;R_j_) for all pairs of the seven classes. Negative values indicate information shared by the two classes; under the conditional-independence model, ΔI_ij_ = −I(R_i_;R_j_) and cannot be positive (Materials and methods). White lines separate basiconic and trichoid classes. Each term was bias-corrected by shuffling as in Fig 2. (B) Interaction information plotted against the Pearson correlation between the mean tuning curves of the two classes (signal correlation). Pairs that include ab2 are shown in orange. Labeled points have ΔI < −0.16 bits.

Redundancy was only weakly related to the similarity of tuning curves (Spearman ρ = −0.34, P = 0.13; Fig 3B). For example, ab1 and ab4 had highly correlated tuning (r = 0.88) but shared less information (−0.15 bits) than at1 and at3 (r = 0.50; −0.28 bits). Shared information therefore depended on which odors the classes resolved reliably, not simply on the overall similarity of their mean profiles.

### A Bayesian decoder identifies 1-octen-3-ol best and reveals confusable odor groups

Mutual information bounds the performance of any decoder but does not say which odors are confused. We therefore constructed a maximum a posteriori decoder that received one response from each of the four basiconic classes on each trial and inferred the odorant under the Gaussian model (Materials and methods). The decoder identified the correct odorant on 32.3% of trials, 5.2 times the chance rate of 6.25% (Fig 4).

**Fig 4.**
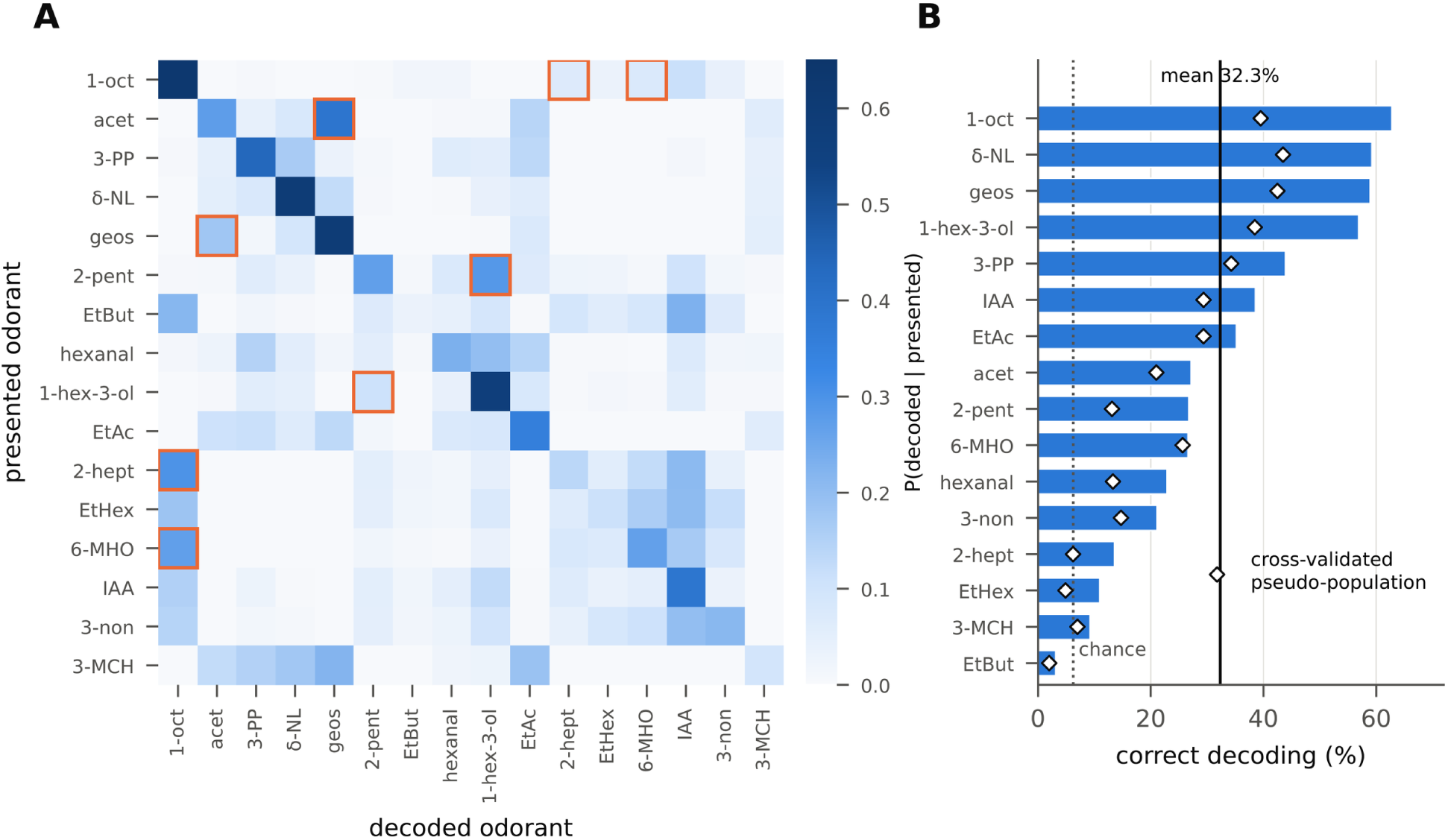
Bayesian decoding of odor identity from the basiconic population. (A) Confusion matrix of a maximum a posteriori decoder that receives one response from each basiconic class per trial (20,000 simulated trials per odorant). Each row gives the probability of each decoded odorant given the presented odorant; the diagonal gives correct identifications. Orange boxes mark the four most frequently confused pairs. (B) Correct decoding for each odorant, sorted. Bars, model decoder; diamonds, decoder trained on two-thirds of the sensilla of each class and tested on held-out sensilla (300 resamplings). Solid line, mean accuracy of the model decoder (32.3%); dotted line, chance (6.25%).

Performance varied 20-fold across odorants. 1-Octen-3-ol was identified best (63%), followed by δ-nonalactone (59%), geosmin (59%), and 1-hexen-3-ol (57%). Ethyl butyrate was identified least often (3%). Errors were structured. The most frequent confusions were acetone with geosmin (28% of trials, averaged over both directions), 2-pentanol with 1-hexen-3-ol (20%), 1-octen-3-ol with 2-heptanone (18%), and 1-octen-3-ol with 6-methyl-5-hepten-2-one (17%). These pairs were close neighbors on the Fisher–Rao manifold (Fig 1A). Confusions involving 1-octen-3-ol were strongly asymmetric. When 1-octen-3-ol was presented, it was mistaken for any single other odorant on at most 11% of trials (isoamyl acetate). Conversely, other strong excitants were often decoded as 1-octen-3-ol, including 2-heptanone (30% of trials), 6-methyl-5-hepten-2-one (27%), and ethyl butyrate (22%). 1-Octen-3-ol thus acted as the default interpretation of strong basiconic activation.

These patterns have plausible ecological correlates. Ethyl butyrate, a fruity ester of fermenting fruit, drove ab3 and ab4 strongly (population d′ = 5.3; see below) but was identified on only 3% of trials: its basiconic response pattern fell within the cluster of strong excitants, and it was decoded as isoamyl acetate (23%) or 1-octen-3-ol (22%). In *Drosophila*, which feeds and oviposits on fermenting fruit, ethyl butyrate strongly activates several dedicated receptors, including Or22a, Or43b, and Or67a [11]. An obligate blood feeder has no evident need to distinguish a fruit ester from other strong volatiles, and the tsetse basiconic channels do not do so; as shown below, however, the trichoid sensilla largely resolved it. The confusion of acetone with geosmin arises because neither odor drives the sampled sensilla strongly at 10^−2^ (5–48 spikes/s in the six classes other than ab2), so both appear as weak activation. This is notable for acetone, a component of host breath and a classic tsetse bait [7,8]. Its behavioral potency may depend on the high release rates used in the field, or on sensilla outside the two antennal regions sampled, which remain to be identified. Geosmin, an earthy microbial volatile, is detected in *Drosophila* by a dedicated aversive channel [29]; no sampled tsetse class plays a comparable role. These interpretations concern what the sampled sensilla can distinguish at a single concentration, not what the fly perceives, and they generate behavioral predictions that can be tested. Because this decoder is built from the same data it is tested on, we also trained a decoder on two-thirds of the sensilla of each class and tested it on pseudo-population responses assembled from the held-out third, repeated 300 times. The cross-validated accuracy was lower (22.8%), as expected when class parameters must be estimated from fewer sensilla, but the per-odorant accuracies of the two decoders were almost perfectly correlated (r = 0.97; Fig 5B). Decoding accuracy was also consistent with the information carried by the fitted model: by Fano’s inequality [30], its 1.40 bits (before bias correction) allow at most 59% correct identification of 16 odors, and the decoder reached 32%.

**Fig 5.**
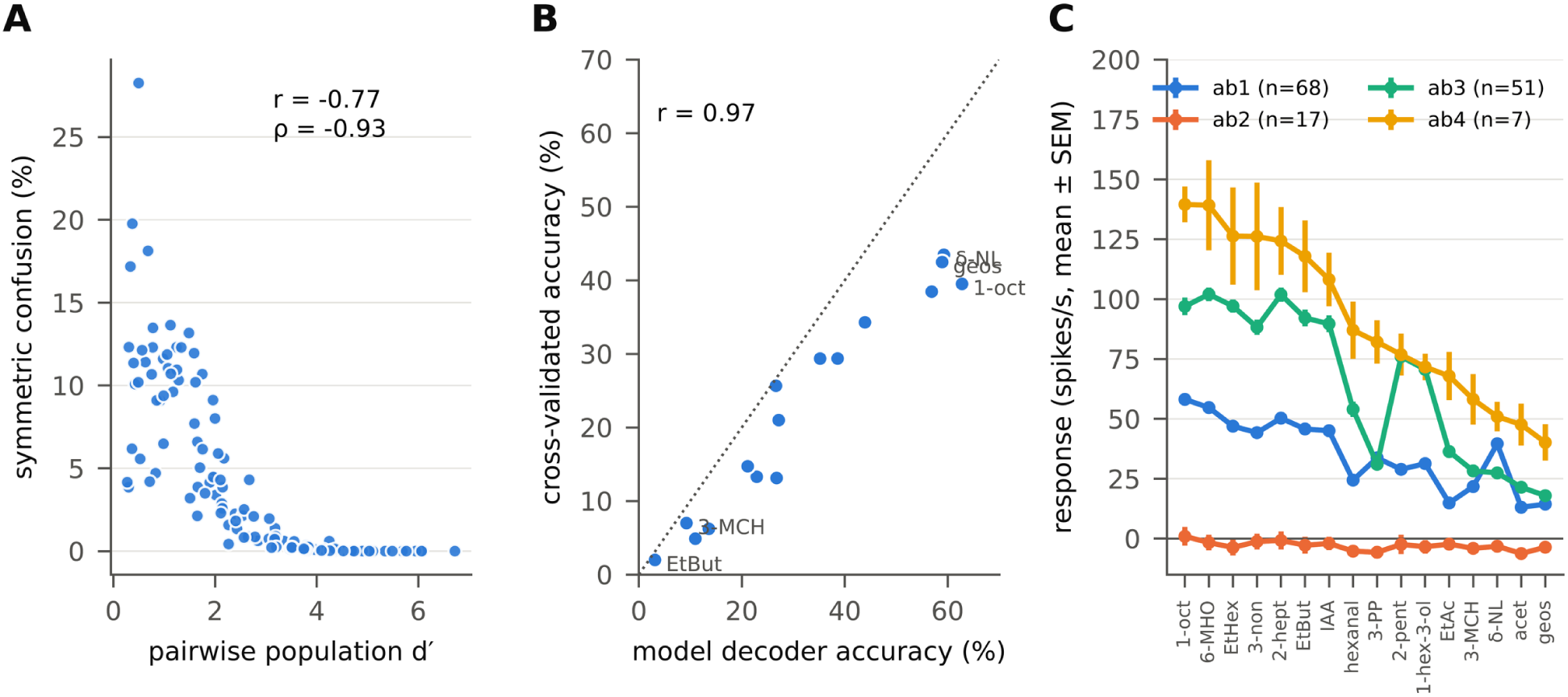
Consistency among analytical frameworks. (A) For each of the 120 pairs of odorants, the symmetric confusion rate of the decoder in Fig 4 plotted against the population d′ separating the pair (Materials and methods). Pearson r = −0.77, Spearman ρ = −0.93 (P < 10^−4^, Mantel permutation test). (B) Per-odorant accuracy of the model decoder compared with that of the cross-validated decoder; dotted line, identity. (C) Mean response profiles (± SEM) of the four basiconic classes, with odorants ordered by the ab4 response. The ab1, ab3, and ab4 profiles are correlated but differ in scale, whereas ab2 responses are small and uniformly negative.

### Detection and discrimination are carried by different classes

Detecting an odor and identifying it are different tasks. Detection asks whether anything is present and requires only a response that is large relative to its variability. Discrimination asks which odor is present and requires responses that differ reliably across odors. A class that fires strongly to every odor can be an excellent detector but a poor discriminator, much as a smoke alarm signals that something is burning without saying what. Mutual information measures discrimination; to measure detection, we used the signal detection index d′ = μ/σ [31,32], the mean baseline-subtracted response of a class divided by its SD across sensilla (Fig 6A). d′ is a signal-to-noise ratio: an observer reading one sensillum drawn at random from a class with d′ = 1 would detect the odor correctly on 69% of presentations, and with d′ > 5 essentially always. Because σ here is the variability across sensilla rather than across repeated trials, d′ measures how consistently a class responds, not the trial-to-trial reliability of one sensillum. In honey bees, behavioral thresholds for detecting odors are likewise lower than those for discriminating them [33]. ab4 had the highest d′ of the four basiconic classes for 11 of 16 odorants (bootstrap 95% range 8–14), including 1-octen-3-ol (d′ = 7.1); ab3 had the highest d′ for the other five. Thus ab4, which carried less information than ab3, was the most sensitive detector, whereas ab3, the most informative, detected fewer odorants best. ab4 responded strongly and consistently enough to signal the presence of most odors, but because it responded strongly to many of them, its response said less about which odor is present. ab4 is also the CO_2_-sensitive class [9]; a single sensillum type may therefore signal two of the most important long-range host cues, CO_2_ and host ketones such as 6-methyl-5-hepten-2-one.

**Fig 6.**
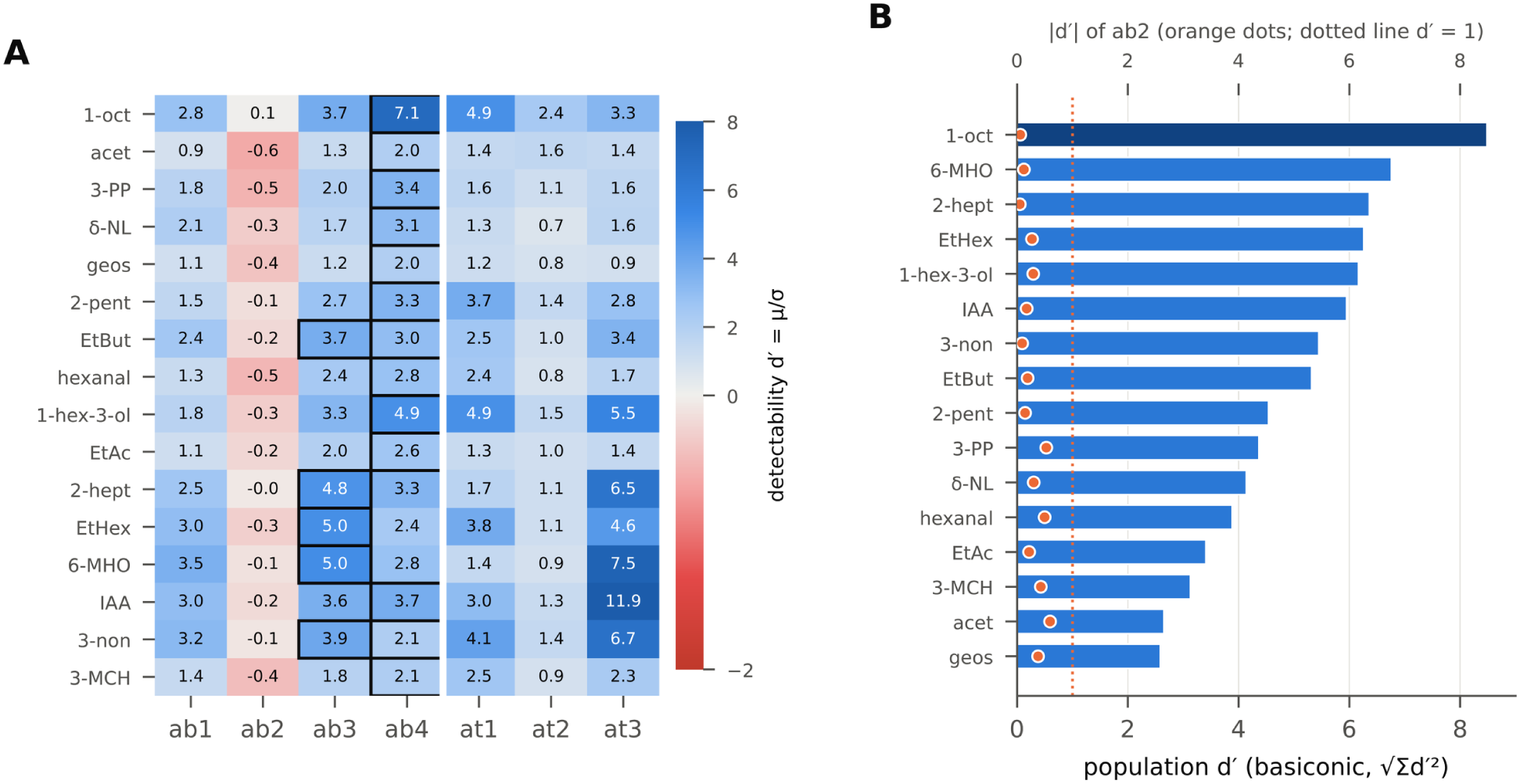
Odor detectability across sensillum classes. (A) Detectability d′ = μ/σ of each odorant for each class, where μ is the mean baseline-subtracted response and σ the SD across sensilla. Black boxes mark the basiconic class with the largest |d′| for each odorant (ab4 for 11 of 16 odorants; ab3 for 5). (B) Population detectability of each odorant by the four basiconic classes, the square root of the summed squared d′ values (bars; bottom axis). Orange dots show |d′| of ab2 alone (top axis); no ab2 response exceeds |d′| = 1 (dotted line).

Combining the four basiconic classes, 1-octen-3-ol was by far the most detectable odorant (population d′ = 8.48), followed by 6-methyl-5-hepten-2-one (6.76) and 2-heptanone (6.36) (Fig 6B). Acetone (2.65) and geosmin (2.59) were the least detectable. Among trichoid classes, at3 detected isoamyl acetate (d′ = 11.9) and 6-methyl-5-hepten-2-one (7.5) especially well, and at1 detected 1-octen-3-ol and 1-hexen-3-ol (d′ = 4.9 each).

ab2 behaved differently from every other class. Its mean response was negative for 15 of 16 odorants, but no response reached |d′| = 1; the largest was the inhibition by acetone (d′ = −0.60). Because these weak inhibitions were nearly uniform across odorants, they conveyed no information about odor identity. Setting all negative ab2 responses to zero did not reveal hidden information (0.035 bits; P = 0.26). An analysis of all 15 combinations of basiconic classes confirmed that ab2 contributed nothing to decoding: adding it changed population information by −0.04 to +0.002 bits, and the small gains in decoding accuracy it produced were matched by ab2 data with shuffled odorant labels (Fig A and Table B in S1 Appendix). In the same analysis, ab4 alone supported the best single-class decoding (19.4%), because it reliably separated strongly from weakly activating odorants.

What might ab2 do? ab2 sensilla have high spontaneous firing rates and appeared to be concentrated near the sensory pit [9]. A neuron with a high baseline can signal in both directions, and odor-evoked inhibition can by itself drive olfactory behavior in *Drosophila* [34]; theory predicts that spontaneous activity combined with inhibitory responses increases coding capacity [22]. If ab2 is tuned to compounds absent from our panel, for example host phenols, pheromone components, or cues associated with larviposition sites, the weak and uniform inhibition observed here would be incidental. Alternatively, it may reflect nonspecific interactions between neurons within the sensillum, such as the ephaptic inhibition described in *Drosophila* [35]. Screening ab2 with broader panels would distinguish these possibilities. For the 16 odorants tested, ab2 was functionally silent, and its removal cost the basiconic code nothing.

Detection and discrimination measures were mutually consistent. Across all 120 odorant pairs, the population d′ separating two odorants predicted how often the decoder confused them (Pearson r = −0.77, Spearman ρ = −0.93; Fig 5A), and Fisher–Rao distance was similarly predictive (r = −0.78). Because pairs that share an odorant are not independent, we assessed significance with a Mantel permutation test, which permutes odorant labels (P < 10^−4^) [36].

### Trichoid sensilla carry most of the information about odor identity

In *Drosophila*, the antennal trichoid sensilla are largely dedicated to pheromones: at1 neurons detect the male pheromone 11-*cis*-vaccenyl acetate [23], and at4 neurons respond to fly-derived odors [24]. Two sensillum types formerly classified as trichoid (at2 and at3), whose receptors respond to some general odorants, have been reclassified as intermediate sensilla [37]. In *G. morsitans*, by contrast, trichoid sensilla on the medial face of the antenna respond to many host odorants [9], and trichoid sensilla on the lateral face contain neurons that respond to a tsetse volatile sex attractant, methyl palmitoleate (MPO), as well as to the host odors 1-octen-3-ol and 4-methylphenol [38]. We therefore asked how much of the tsetse antenna’s information about odor identity is carried by its trichoid sensilla.

The three trichoid classes together carried 1.92 bits about the 16 odorants, more than the four basiconic classes (1.14 bits), and a decoder given one response from each trichoid class identified the correct odor on 52.9% of trials, compared with 32.2% for the basiconic classes (Fig 7A). All seven classes together carried 2.27 bits and were decoded correctly on 69.5% of trials. The trichoid advantage was robust. The difference between the trichoid and basiconic classes, 0.78 bits, had a bootstrap 95% CI of 0.44–1.10 bits, and no bootstrap replicate reversed it. Decoders tested on held-out sensilla were less accurate, as expected, but kept the same order: 22.8% for the basiconic classes, 39.7% for the trichoid classes, and 54.6% for all seven (Table F in S1 Appendix). In simulations at the real sample sizes, the shuffle correction underestimated trichoid information more than basiconic information, so the difference is, if anything, conservative (Table E in S1 Appendix). Finally, because the classes were defined by clustering the same responses, we asked whether this circularity inflates information. When classes were defined by clustering on half of the odorants and information was measured on the other half, basiconic information fell by 8% and trichoid information did not fall (Table E in S1 Appendix), so circular class definition does not explain the trichoid advantage. The overlap between the two sensillum types was asymmetric. Adding the trichoid classes to the basiconic classes added 1.13 bits, whereas adding the basiconic classes to the trichoid classes added only 0.35 bits. Most of what the basiconic population conveyed about odor identity was therefore also conveyed by the trichoid population, but not the reverse.

**Fig 7.**
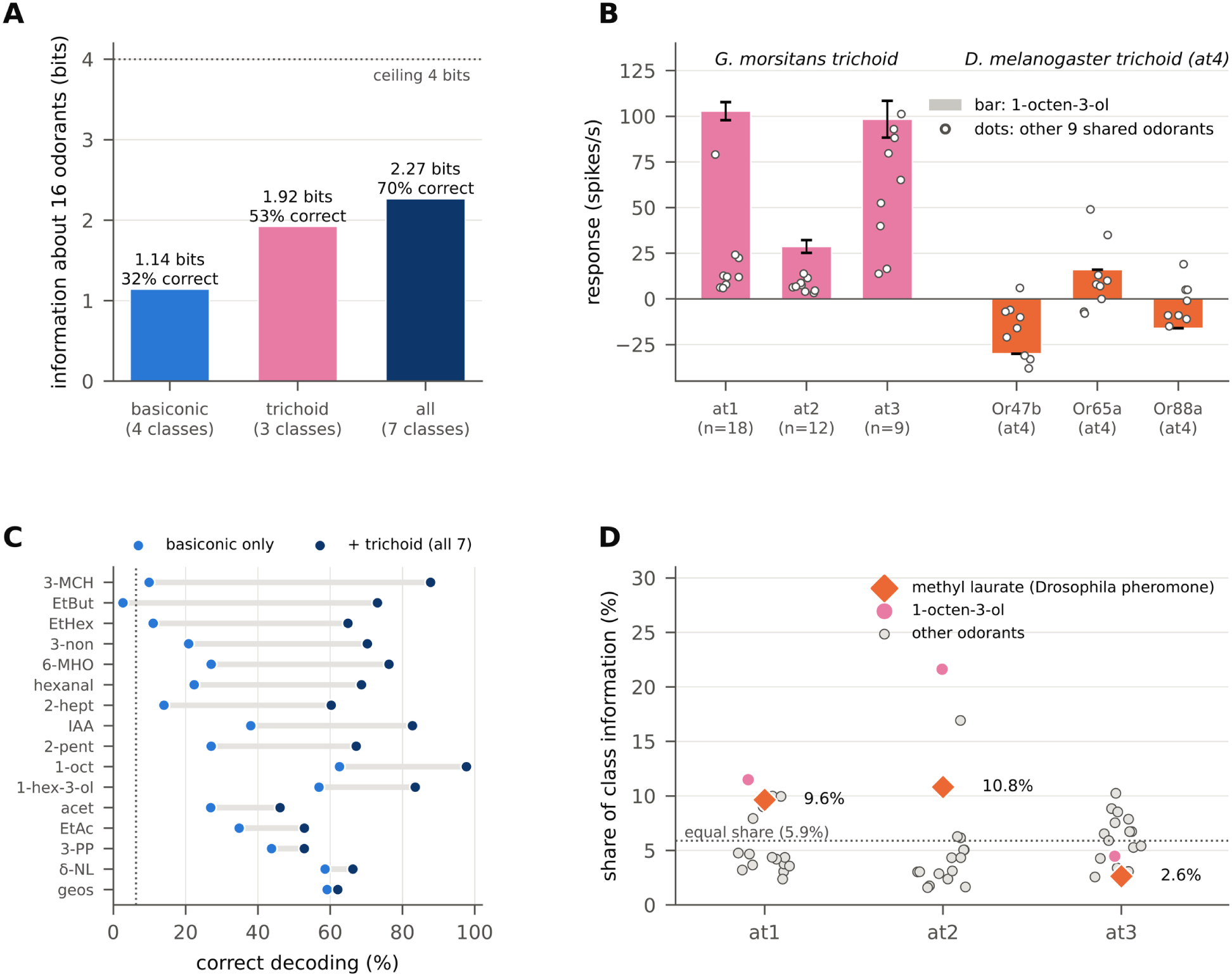
Trichoid sensilla carry most of the information about odor identity in the tsetse antenna. (A) Information about the identity of 16 odorants carried by one response from each basiconic class, each trichoid class, or all seven classes (bias-corrected), with the corresponding accuracy of the Bayesian decoder. Dotted line, ceiling of 4 bits. (B) Responses of trichoid channels to 1-octen-3-ol (bars) and to the nine other odorants tested in both species (dots). Left, *G. morsitans* trichoid classes (mean ± standard error of the mean, SEM). Right, *Drosophila* at4 trichoid receptors expressed in the “empty neuron” (data from [11]). (C) Correct decoding of each odorant from the basiconic classes alone (blue) and from all seven classes (dark blue), sorted by the gain. Dotted line, chance (6.25%). (D) Share of each trichoid class’s information about 17 odorants contributed by each odorant (specific surprise). Diamonds, methyl laurate, a *Drosophila* pheromone; pink, 1-octen-3-ol; gray, other odorants. Dotted line, the share expected if all odorants contributed equally (5.9%).

Trichoid sensilla were strongly driven by the host attractant 1-octen-3-ol. at1 and at3 responded with about 100 spikes/s (102.8 ± 4.9 and 98.3 ± 10.0 spikes/s), and 1-octen-3-ol was the strongest stimulus for at1 and at2 (Fig 7B). The *Drosophila* at4 receptors, in contrast, were not excited by 1-octen-3-ol (Or47b, −30; Or65a, 16; Or88a, −16 spikes/s) and responded weakly or not at all to the other nine odorants shared between the two panels (maximum 49 spikes/s, Or65a to 6-methyl-5-hepten-2-one) [11]. The same morphological type of sensillum that in *Drosophila* is largely insensitive to food and host volatiles was, in tsetse, one of the principal detectors of a key host odor.

The information added by the trichoid classes was concentrated on the odors that the basiconic classes could not tell apart (Fig 7C; Table D in S1 Appendix). Identification of 3-methyl-2-cyclohexenone rose from 10% to 88%, ethyl butyrate from 3% to 73%, ethyl hexanoate from 11% to 65%, 6-methyl-5-hepten-2-one from 27% to 76%, and 2-heptanone from 14% to 60%. Much of this gain came from at3, which detected isoamyl acetate (d′ = 11.9), 6-methyl-5-hepten-2-one (7.5), and 2-heptanone (6.5) with a tuning profile distinct from those of the basiconic classes. 1-Octen-3-ol was identified on 98% of trials (92% with held-out sensilla), and its confusions with 2-heptanone and 6-methyl-5-hepten-2-one, 18% and 17% with basiconic sensilla alone, fell to zero. Some confusions persisted: acetone with geosmin (25%), which no class encoded strongly; 2-pentanol with 1-hexen-3-ol (22%), two alcohols that drove at1 and the basiconic classes alike; ethyl hexanoate with 3-nonanone (19%); and the two host ketones 2-heptanone and 6-methyl-5-hepten-2-one (17%). Conversely, basiconic sensilla were the better channels for 3-propylphenol, δ-nonalactone, and geosmin, which the trichoid classes alone identified on only 18%, 30%, and 53% of trials, compared with 44%, 59%, and 59% for the basiconic classes. The two sensillum types thus divided the labor: basiconic sensilla signaled the presence and strength of host odors and resolved the phenol, the lactone repellent, and weak odorants, whereas trichoid sensilla separated strong host ketones and esters from 1-octen-3-ol and from many other odorants, although some pairs, such as 2-heptanone and 6-methyl-5-hepten-2-one, remained partly confusable.

Finally, we asked how much of the trichoid information about the panel concerns methyl laurate, the one fly-derived compound tested. Methyl laurate is a *Drosophila* pheromone, detected by at4 neurons expressing Or47b and Or88a [39], the same trichoid receptors that are insensitive to the host and food odors in Fig 7B; it is not known to act as a pheromone in tsetse. We decomposed the information carried by each trichoid class about all 17 odorants into the contribution of each odorant, using the specific surprise [40]. If every odorant contributed equally, each would account for 5.9%. Methyl laurate accounted for 9.6% of the information carried by at1, 10.8% for at2, and 2.6% for at3 (Fig 7D). In at1 and at2, 1-octen-3-ol contributed more than methyl laurate (11.5% and 21.6%). Even in the classes that respond best to methyl laurate, therefore, this *Drosophila* pheromone contributed a share comparable to that of single host odorants. These shares describe the tested panel only. The panel did not include the tsetse volatile sex attractant MPO or the related methyl esters, which excite trichoid neurons on the lateral face of the antenna [38]. The present data therefore show that, on the medial face sampled here, host odors account for most of the trichoid information about this panel; they do not measure how tsetse trichoid sensilla divide their capacity between host odors and tsetse pheromones.

### The tsetse antenna carries less odor information than the *Drosophila* antenna

Finally, we asked whether these properties distinguish tsetse from *Drosophila*. Ten odorants were tested at the same 10^−2^ dilution both in *G. morsitans* and in the comprehensive survey of 24 *Drosophila* antennal odorant receptors by Hallem and Carlson [11] (Fig 8A). Because the *Drosophila* data are reported as means, we fitted a noise model to the tsetse data, in which the SD across sensilla grows linearly with response magnitude (σ = 7.7 + 0.20|μ| spikes/s), and applied the same model to both species. This comparison therefore isolates differences in tuning from differences in variability.

**Fig 8.**
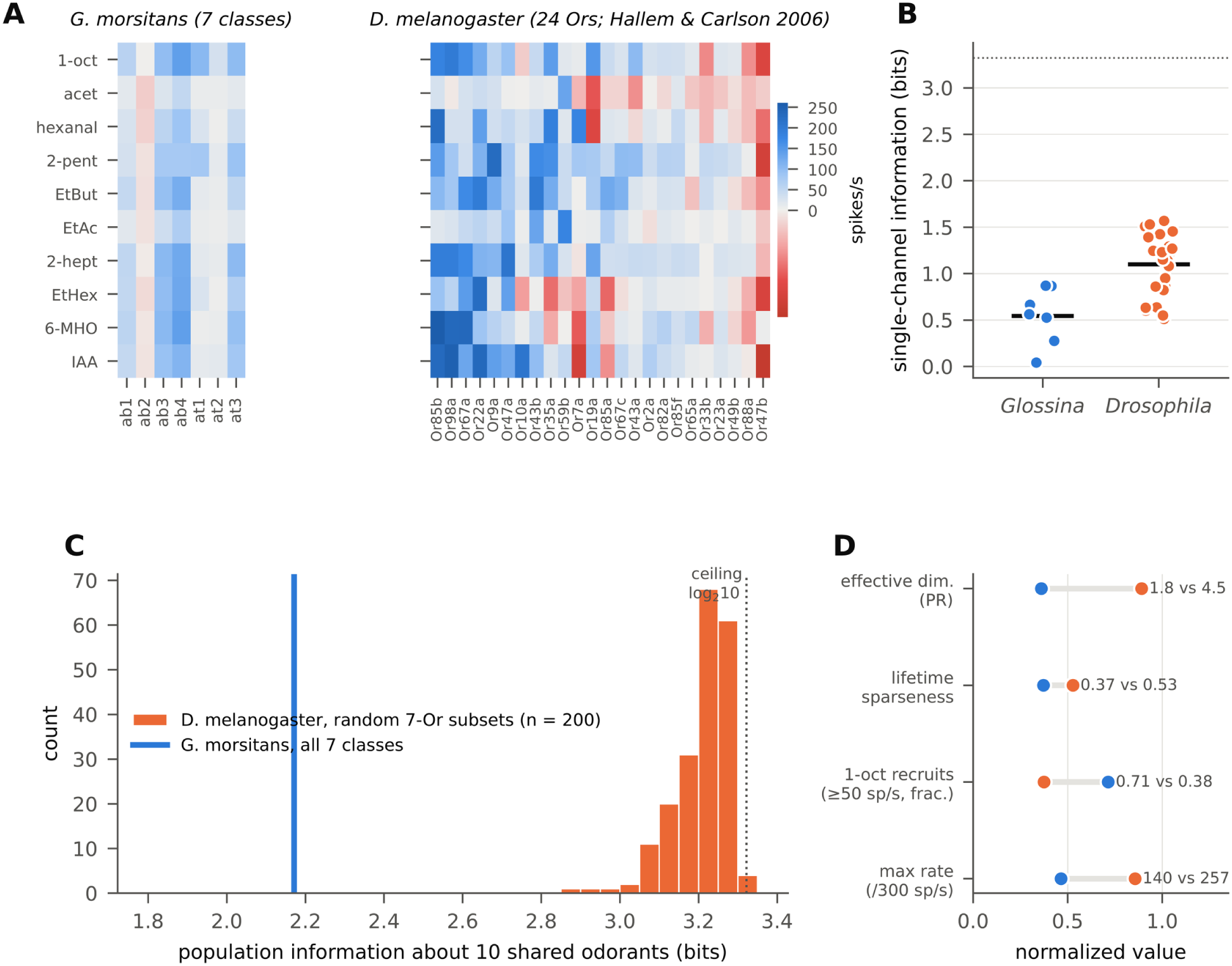
Comparison of odor coding in *G. morsitans* and *D. melanogaster* on 10 odorants tested at 10^−2^ in both species. (A) Mean responses of the seven *G. morsitans* sensillum classes (left) and of 24 *Drosophila* odorant receptors expressed in the “empty neuron” system (right; data from [11]), on a common color scale (spikes/s). *Drosophila* receptors are ordered by their maximal response. (B) Information carried by each single channel about the 10 odorants, computed with the same noise model for both species (σ = 7.7 + 0.20|μ| spikes/s, fitted to the tsetse data). Horizontal bars, means. Dotted line, ceiling (log_2_10). (C) Population information for random sets of 7 *Drosophila* receptors (histogram; 200 sets) and for all 7 *G. morsitans* classes (blue line). (D) Summary of coding properties, each normalized for display: effective dimensionality (participation ratio across all channels, divided by 5; not matched for channel number, see text); mean lifetime sparseness; fraction of channels excited by 1-octen-3-ol at ≥ 50 spikes/s; and maximal response (divided by 300 spikes/s). Blue, *G. morsitans*; orange, *Drosophila*.

Under matched noise, a single *Drosophila* receptor carried on average twice as much information about the 10 odorants as a single *G. morsitans* sensillum class (1.10 vs. 0.54 bits; Fig 8B; Table C in S1 Appendix).

To control for the number of channels, we drew 200 random sets of 7 *Drosophila* receptors, matching the seven *G. morsitans* classes. These sets carried a median of 3.23 bits (95% range 3.06–3.30), close to the 3.32-bit ceiling, compared with 2.17 bits for the seven *G. morsitans* classes; none of the 200 *Drosophila* sets carried as little information as *G. morsitans* (Fig 8C). The *Drosophila* receptors spanned more dimensions of odor space (participation ratio 4.46 vs. 1.80 across channels). Because the participation ratio grows with the number of channels, we also computed it for random sets of 7 *Drosophila* receptors: these gave a median of 2.79 (95% range 1.79–3.87), and only 2.7% of sets were as low-dimensional as the *G. morsitans* classes. The *Drosophila* receptors were also more selective (mean lifetime sparseness 0.53 vs. 0.37), and reached higher firing rates (maximum 257 vs. 140 spikes/s on this panel) (Fig 8D). In contrast, 1-octen-3-ol elicited ≥ 50 spikes/s from 5 of 7 *G. morsitans* classes (71%) but from only 9 of 24 *Drosophila* receptors (38%). The information difference between species held under constant, proportional, and doubled noise models (Table G in S1 Appendix).

## Discussion

By asking how much information the tsetse antenna carries rather than how its sensilla cluster, this reanalysis identifies several organizing principles of the tsetse peripheral olfactory code. The most striking concerns the trichoid sensilla. The basiconic code is nearly one-dimensional, redundant, and highly sensitive to host odors, but it is the trichoid sensilla that carry most of the information about which host odor is present.

### Trichoid sensilla as host-odor channels

In *Drosophila*, trichoid sensilla are a pheromone-detecting system: their neurons detect fly-derived compounds such as 11-*cis*-vaccenyl acetate [23,24], and the at4 receptors are not excited by the host and food volatiles shared with our panel [11]. The original survey showed that tsetse trichoid sensilla respond to many host odors [9]. The present analysis shows that this is not a minor addition to the tsetse code but, for the panel tested, its main source of odor identity: the three trichoid classes carry more information than the four basiconic classes, contain most of the basiconic information, and resolve many confusions among host ketones and esters that the basiconic classes cannot. Tsetse trichoid sensilla are not, however, devoid of pheromone function. Ebrahim et al. identified MPO as a volatile sex attractant of *G. morsitans* and found that the trichoid neurons it excites also respond strongly to 1-octen-3-ol and 4-methylphenol [38]. Together, the two studies suggest that tsetse trichoid sensilla are dual-purpose channels for host odors and pheromones, rather than the largely pheromone-dedicated system of *Drosophila*. The same morphological sensillum type thus appears to serve different ecological functions in the two species. *Drosophila* uses olfaction to find mates, food, and oviposition sites among a wide range of fermenting substrates. Tsetse feed exclusively on vertebrate blood, and females bear live larvae rather than laying eggs. A shift of trichoid sensilla toward host-odor identification is consistent with this ecology. Whether it reflects a change in the receptors expressed in trichoid neurons, which could be tested by mapping tsetse receptor expression to sensillum types [41], remains to be determined.

### Detection versus discrimination

Signal detection and information analyses assigned distinct roles to the sensillum classes. Among basiconic classes, ab4 was the most sensitive detector for most odorants, whereas ab3 carried the most information about odor identity; across the antenna, basiconic sensilla signal the presence and strength of host odors and trichoid sensilla add their identity. These roles need not conflict. A channel with large responses to many odors, such as ab4, reliably signals that a strong stimulus is present but distinguishes poorly among strong stimuli, whereas channels with more differentiated profiles resolve identity. For a fly flying upwind in an intermittent odor plume, detecting a host signal quickly and reliably may be as important as identifying its source, and the antenna appears to allocate separate channels to the two tasks. ab4 is also the CO_2_-sensitive class [9], so a single sensillum type may integrate two of the most important long-range host cues, CO_2_ and ketones such as 6-methyl-5-hepten-2-one. The dominance of a response-magnitude axis in the Fisher–Rao geometry suggests that a large part of what the tsetse antenna conveys is how strongly an odor activates the host-odor channels as a whole, a quantity well suited to guiding upwind flight toward a source. An alternative account deserves testing: all odorants were delivered at a 10^−2^ dilution, and odorants differ widely in volatility, so part of this axis may reflect how much of each odorant reached the antenna. Recordings at concentrations matched for vapor-phase delivery would separate the two accounts.

### The geometry of 1-octen-3-ol

1-Octen-3-ol was the most detectable odorant, the best-decoded odorant, and occupied the strong-response pole of the odor manifold. Earlier recordings from several *Glossina* species identified cells sensitive to 1-octen-3-ol [42,43], and field studies established it as a potent tsetse attractant [7,8]. Our analysis adds a quantitative rationale: because 1-octen-3-ol drives nearly every excitatory channel strongly, it is maximally separated from background and from most other odors. In the basiconic population alone, other strong excitants were frequently decoded as 1-octen-3-ol, whereas the reverse error was rare, so that 1-octen-3-ol behaves as the strongest instance of a general “host” signal. The trichoid sensilla remove this ambiguity: with all seven classes, 1-octen-3-ol was identified on 98% of trials. Tsetse may therefore represent 1-octen-3-ol both as an intense, readily detected signal through basiconic sensilla and as a specific identity through trichoid sensilla.

### Redundancy as a design choice

Thirty-two percent of basiconic information and 45% of the information across all seven classes was shared between classes; simulations indicate that these fractions are upper estimates (Table E in S1 Appendix). Because classes were combined under conditional independence, this analysis can detect only redundancy arising from shared tuning; correlated variability between classes, which simultaneous recordings could measure, could either add or remove information. Redundancy lowers capacity but increases robustness: information about host odor survives the loss or adaptation of any single class and can be averaged across the many sensilla of each class to overcome noise [44]. Because we quantified information per sensillum, with sensillum-to-sensillum variability as noise, pooling across the hundreds of sensilla of each class would increase the information available to the brain, and correlated class profiles make that pooling effective. The shared profiles of ab1, ab3, ab4, and at3 are consistent with the previously proposed sharing of an odorant receptor neuron or receptor among sensillum classes [9], an arrangement that would naturally produce this redundancy. For a blood-feeding fly that must track a sparse, intermittent host plume, robust detection may be worth more than fine discrimination.

### Reinterpreting ab2

ab2 was the one class without excitatory responses in the original survey [9], and our analyses detected no information about this panel in its responses (upper 95% confidence bound, 0.07 bits). As discussed above, ab2 may be tuned to compounds outside the panel, in which case its high spontaneous activity would allow bidirectional coding once its ligands are found [22,34], or its weak, uniform inhibition may reflect interactions within the sensillum [35]. Identifying the ligands of ab2 and at2, the two classes without strong excitatory responses, is a clear priority for completing the functional map of the tsetse antenna.

### Divergence from *Drosophila*

On a common panel and under matched noise, *Drosophila* receptors carried more information, spanned more dimensions even when channel number was matched, and were more selective than *G. morsitans* sensillum classes, whereas 1-octen-3-ol recruited a larger fraction of *G. morsitans* channels. This difference holds even though tsetse trichoid sensilla add substantially to the tsetse code. The *Drosophila* antenna must represent the chemically diverse volatiles of fermenting fruit and suitable oviposition sites [11,20]; the tsetse antenna serves an obligate blood feeder whose larvae develop in utero. A peripheral code dominated by a single host-odor axis is consistent with this narrower ecological demand. The comparison should be interpreted with caution, because the *Drosophila* data derive from single receptors expressed in a common neuron and the *G. morsitans* data from summed activity of all neurons in a sensillum. Summation may blur the tuning of individual neurons, so single tsetse neurons may be more selective than the totals of their sensilla suggest.

### Implications for odor baits

These results suggest principles for designing tsetse attractants. First, because the basiconic code is dominated by overall activation of the host-odor channels, blends that drive ab3, ab4, at1, and at3 strongly are expected to be most salient. Second, because the identity of host odors is carried largely by trichoid sensilla, candidate attractants should be screened on trichoid as well as basiconic sensilla, and synthetic blends that mimic a host should reproduce the trichoid response pattern as well as the overall intensity. Third, odors that remain confusable across all seven classes, such as acetone with geosmin, 2-pentanol with 1-hexen-3-ol, or 2-heptanone with 6-methyl-5-hepten-2-one, may be partially interchangeable in baits, which could be tested behaviorally. Fourth, the decoding framework can predict whether a candidate repellent, such as δ-nonalactone, is represented distinctly from attractants; notably, δ-nonalactone was among the best-decoded odorants and had the largest nearest-neighbor distance of any odorant on the manifold (d = 1.34).

### Limitations

This analysis has several limitations. First, responses were summed across all neurons in a sensillum, so our estimates describe sensillum classes rather than individual receptor neurons. Second, the classes were defined by clustering the same responses, which may make within-class responses more homogeneous than they would be under an independent classification; a split-half test suggested that this inflates basiconic information by about 8% and trichoid information not at all. Third, variability was measured across sensilla, not across repeated trials, and joint information assumed that responses of different classes are conditionally independent given the odor; each is a reasonable approximation for classes housed in different sensilla, but conditional independence means that synergy between classes could not be detected, and it should be tested with simultaneous recordings. Fourth, all odorants were tested at a single dilution, so information about concentration was not addressed and differences in volatility may contribute to the response-magnitude axis. Recordings also sampled particular antennal regions, mainly around the sensory pit, with ab4 recorded more distally [9], so classes elsewhere on the antenna may code odors differently. Fifth, ab4 (n = 7) and at3 (n = 9) were sampled sparsely, as reflected in their confidence intervals. Sixth, the *Drosophila* trichoid comparison relies on receptors expressed in a heterologous neuron, and the at1 receptor Or67d is not included in that dataset. Finally, the cross-species comparison relied on a noise model fitted to tsetse; because this model assigns small variability to weak responses, it may overestimate the information carried by weakly responding *Drosophila* receptors, so for the trichoid comparison we emphasize response magnitudes. The four frameworks share one response model, so their agreement (Fig 5) is a check of internal consistency rather than independent validation. The main conclusions did, however, hold for decoders tested on held-out sensilla, in simulations at the real sample sizes, under alternative noise models, and when classes were defined on independent odorants (Tables E–G in S1 Appendix).

In summary, the tsetse antenna combines a low-dimensional, redundant, detection-oriented basiconic code, in which host odors and 1-octen-3-ol above all stand out, with trichoid sensilla that carry most of the information about host-odor identity in the panel tested, alongside their newly identified role in pheromone detection, a combination quite different from the pheromone-dedicated trichoid sensilla of *Drosophila*. Quantifying information rather than describing clusters provides a framework for comparing olfactory systems across insects and for rationally designing the odor baits used to control disease vectors.

## Materials and methods

### Data

All electrophysiological data were published previously [9], and experimental procedures are described there. Briefly, single-sensillum recordings were made from the antennae of *G. morsitans morsitans*; each odorant was diluted to 10^−2^ in paraffin oil (methyl laurate was used neat) and delivered as a 500-ms pulse, and responses were quantified as the summed spike frequency of all neurons in the sensillum over 500 ms, with the spontaneous rate and the response to the paraffin oil diluent subtracted. We analyzed 143 basiconic sensilla (ab1, n = 68; ab2, n = 17; ab3, n = 51; ab4, n = 7), each tested with 16 odorants, and 39 trichoid sensilla (at1, n = 18; at2, n = 12; at3, n = 9), each tested with the same 16 odorants and methyl laurate. Class assignments were those of the original study; mean responses reproduced the published values [9]. Analyses used Python 3.13 with NumPy 2.5, SciPy 1.18, pandas 3.0, scikit-learn 1.9, matplotlib 3.11, and drosolf 0.1.3; all random number generators were seeded. No new experiments were performed for this study, and work on insects does not require institutional animal care approval.

### Response model

For class c and odorant s, responses across sensilla were modeled as Gaussian, R_c_ | s ∼ N(μ_sc_, σ_sc_^2^), with μ_sc_ and σ_sc_ the sample mean and SD (floor 1 spike/s). Responses of different classes were assumed conditionally independent given the odorant.

### Information geometry

The Fisher–Rao distance between two univariate Gaussians has the closed form [45]

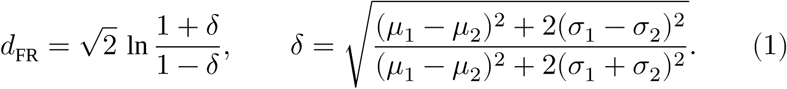

Because the population distribution is a product over classes, the distance between odorants i and j on the product manifold is [16]

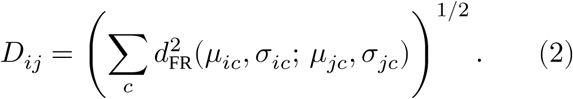

The matrix D was embedded by classical multidimensional scaling [46]; the fraction of geometry on each axis is the corresponding positive eigenvalue of the double-centered matrix −½J*D*^∘2^J divided by the sum of positive eigenvalues, where J is the centering matrix and *D*^∘2^ the element-wise square of D. Distance to the centroid was computed in the full embedding. Effective dimensionality was measured by the participation ratio [25]

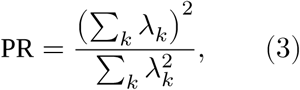

where λ_k_ are the eigenvalues of the response covariance matrix.

### Mutual information

With S uniform over N_s_ odorants, the mutual information between odorant and response is [12,30]

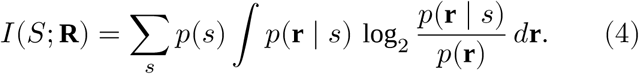

For single classes the integral was evaluated numerically on a 4,000-point grid; for multiple classes it was estimated by Monte Carlo integration (4 × 10^4^ samples). Finite samples bias plug-in estimates upward [26,47].

We estimated this bias by independently permuting the odorant labels within each sensillum, which preserves each sensillum’s response distribution while destroying odorant identity, recomputing the information, and subtracting the mean over 100 permutations (40 for combinations of classes). The permutation distribution also provided P values. Confidence intervals were ±1.96 bootstrap SE, obtained by resampling sensilla with replacement within each class (200 replicates for single classes, 100 for the basiconic population), each replicate corrected by its own shuffle null (10 and 5 permutations, respectively). As a distribution-free check, we computed the k-nearest-neighbor estimator for mixed discrete–continuous variables (k = 3) [27,28]. Interaction information for classes i and j was computed from bias-corrected terms as [14]

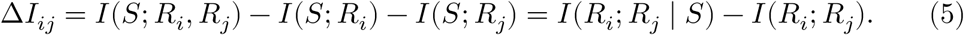

Because responses of different classes were modeled as conditionally independent given the odorant, I(R_i_;R_j_ | S) = 0 and ΔI_ij_ = −I(R_i_;R_j_) ≤ 0. In this model ΔI therefore quantifies information shared through tuning (signal redundancy) and cannot reveal synergy, which would require correlated variability between classes. We verified the identity numerically for four pairs by computing I(R_i_;R_j_) directly (maximum discrepancy 0.003 bits).

Redundancy was defined as Σ_c_ I(S;R_c_) − I(S;**R**). We report information for equally likely odorants in our panel; this is a lower bound on the channel capacity for that panel, which is the maximum over all odorant distributions. Stimulus-specific contributions were computed as the specific surprise [40]

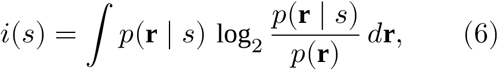

which averages to I(S;**R**); the share contributed by each odorant is i(s)/Σ_s′_ i(s′).

### Bayesian decoding

On each simulated trial an odorant s was drawn, a response r_c_ was drawn from N(μ_sc_, σ_sc_^2^) for each class in the decoding set, and the decoder chose the maximum a posteriori odorant under a uniform prior,

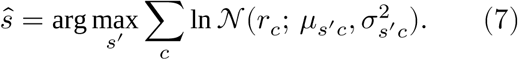

We simulated 20,000 trials per odorant for the full basiconic set and 6,000 for each of the 15 combinations of basiconic classes, for the trichoid classes, and for all seven classes. As a consistency check, decoding accuracy was compared with the upper limit set by Fano’s inequality [30], using the information of the fitted model before bias correction, because the decoder uses the fitted parameters,

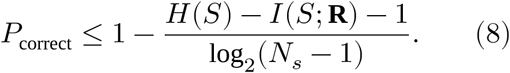

To test whether small gains in accuracy from adding ab2 exceeded chance, we repeated the decoding with ab2 data whose odorant labels were shuffled within each sensillum (20 shuffles for ab2 alone, 10 for the four-class set). For cross-validation, one-third of the sensilla of each class (at least one) were held out; the decoder parameters were estimated from the remainder, and test trials were pseudo-population vectors formed by drawing one held-out sensillum of each class (40 trials per odorant per split; 300 splits).

### Signal detection

Detectability was d′_sc_ = μ_sc_/σ_sc_, the baseline- and diluent-subtracted response in units of its SD [31,32]. For an unbiased observer deciding between odor and diluent from one response, the probability of a correct decision is Φ(d′/2), where Φ is the standard normal cumulative distribution function. Population detectability was (Σ_c_ d′_sc_^2^)^1/2^. Pairwise discriminability of odorants i and j was

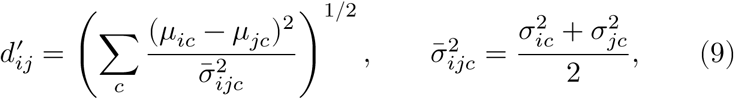

and was compared with the symmetric confusion rate (C_ij_ + C_ji_)/2 of the decoder.

### Cross-species comparison

*Drosophila* responses of 24 odorant receptors expressed in the ab3A “empty neuron” [11] were obtained from the published table (via the drosolf Python package). Ten odorants were common to both panels at 10^−2^: 1-octen-3-ol, acetone, hexanal, 2-pentanol, ethyl butyrate, ethyl acetate, 2-heptanone, ethyl hexanoate, 6-methyl-5-hepten-2-one, and isoamyl (isopentyl) acetate. A linear noise model, σ = a + b|μ|, was fitted by least squares to all odorant–class pairs of the seven tsetse classes (a = 7.7 spikes/s, b = 0.20) and applied to both species. Lifetime sparseness was computed on rectified responses [48]. Channel-matched comparisons used 200 random sets of 7 *Drosophila* receptors. For the trichoid comparison, receptors were assigned to sensillum types according to [37,49]; the at4 receptors Or47b, Or65a, and Or88a are the *Drosophila* trichoid receptors represented in the dataset.

### Robustness and validation analyses

Parametric recovery: for single classes (ab3, ab4, at2, at3) and for the basiconic, trichoid, and seven-class populations, we treated the fitted means and SDs as ground truth, simulated datasets with the real number of sensilla per class (40 replicates for single classes, 25 for populations), and applied the full estimator (plug-in information minus a 15-permutation shuffle null). The bias of the estimator is the mean estimate minus the true information of the fitted model. The trichoid–basiconic difference was bootstrapped by resampling sensilla within each class (150 replicates, each corrected by its own 5-permutation shuffle null); the CI is the 2.5th–97.5th percentile. Held-out decoding for the trichoid and seven-class sets used the cross-validation procedure described above. The stability of the best-detector assignment was assessed by recomputing d′ on 1,000 bootstrap resamples of sensilla. The correlation between pairwise d′ and decoder confusion was tested with a Mantel test, permuting odorant labels of the d′ matri× 10,000 times [36]. For the cross-species comparison, the participation ratio was recomputed for 1,000 random sets of 7 *Drosophila* receptors, and per-channel and population information were recomputed under three further noise models: constant (σ = 20 spikes/s), proportional (σ = 1 + 0.3|μ|), and the fitted model with both coefficients doubled. To estimate how much defining classes on the same data inflates information, we split the 16 odorants at random into two halves (30 splits), clustered the basiconic or trichoid sensilla into four or three groups by Ward’s method [50] on one half, and computed the population information on the other half; this was compared with clustering and measuring on the same half. All these analyses are in the script robustness.py.

### Use of AI tools

Claude (Anthropic) was used to write and run the Python analysis and plotting code, to draft the manuscript text and S1 Text under the author’s direction, and to check the internal consistency of the reported numbers. The author conceived the research, designed the study, collected the original data, chose the mathematical and analytical frameworks, interpreted the results, and reviewed and edited all text. All analyses were verified by the author: every reported value is regenerated by the deposited code from the raw data, estimator bias was assessed by simulation with known ground truth (S1 Appendix, Table E), and the author takes full responsibility for the content.

## Data and code availability

The electrophysiological data analyzed here were published previously [9]. The sensillum-by-odorant response matrices and all analysis code, which reproduce every number and figure in this paper from the raw data, are available on GitHub (https://github.com/cellularphysiology/tsetse-information-coding) and archived in Zenodo (version 1.0.0; https://doi.org/10.5281/zenodo.23420875).

## Acknowledgments

I thank John R. Carlson, in whose laboratory the original data were collected.

## Supporting information

**S1 Text. A guide to the mathematical framework.** What each of the four analyses (signal detection theory, Fisher–Rao information geometry, mutual information, and Bayesian decoding) measures, why it is needed, the formulas used, worked examples from the tsetse data, how the frameworks check one another, and the assumptions of the analysis.

**S1 Appendix. Supplementary tables and figure.** Table A, mutual information estimates for each sensillum class. Table B, information and decoding accuracy for all combinations of basiconic classes. Table C, cross-species comparison on 10 shared odorants. Table D, decoding of each odorant by basiconic, trichoid, and all sensillum classes. Fig A, contribution of each basiconic class to decoding and information.

## Notes

### Competing Interest Statement

The authors have declared no competing interest.

## References

1. Simarro PP, Cecchi G, Franco JR, Paone M, Diarra A, Ruiz-Postigo JA, et al. Estimating and mapping the population at risk of sleeping sickness. PLoS Negl Trop Dis. 2012;6: e1859.

2. Budd LT. DFID-funded tsetse and trypanosome research and development since 1980. Vol. 2: Economic analysis. London: Department for International Development; 1999.

3. Franco JR, Priotto G, Paone M, Cecchi G, Ebeja AK, Simarro PP, et al. The elimination of human African trypanosomiasis: Monitoring progress towards the 2021–2030 WHO road map targets. PLoS Negl Trop Dis. 2024;18: e0012111.

4. Lejon V, Lindner AK, Franco JR. Human African trypanosomiasis. Lancet. 2025;405: 937–950.

5. Tirados I, Esterhuizen J, Kovacic V, Mangwiro TN, Vale GA, Hastings I, et al. Tsetse control and Gambian sleeping sickness; implications for control strategy. PLoS Negl Trop Dis. 2015;9: e0003822.

6. Gibson G, Torr SJ. Visual and olfactory responses of haematophagous Diptera to host stimuli. Med Vet Entomol. 1999;13: 2–23.

7. Torr SJ. Dose responses of tsetse flies (*Glossina*) to carbon dioxide, acetone and octenol in the field. Physiol Entomol. 1990;15: 93–103.

8. Vale GA, Hall DR. The use of 1-octen-3-ol, acetone and carbon dioxide to improve baits for tsetse flies, *Glossina* spp. (Diptera: Glossinidae). Bull Entomol Res. 1985;75: 219–232.

9. Soni N, Chahda JS, Carlson JR. Odor coding in the antenna of the tsetse fly *Glossina morsitans*. Proc Natl Acad Sci U S A. 2019;116: 14300–14308.

10. de Bruyne M, Foster K, Carlson JR. Odor coding in the *Drosophila* antenna. Neuron. 2001;30: 537–552.

11. Hallem EA, Carlson JR. Coding of odors by a receptor repertoire. Cell. 2006;125: 143–160.

12. Shannon CE. A mathematical theory of communication. Bell Syst Tech J. 1948;27: 379–423.

13. Borst A, Theunissen FE. Information theory and neural coding. Nat Neurosci. 1999;2: 947–957.

14. Schneidman E, Bialek W, Berry MJ 2nd. Synergy, redundancy, and independence in population codes. J Neurosci. 2003;23: 11539–11553.

15. Rao CR. Information and the accuracy attainable in the estimation of statistical parameters. Bull Calcutta Math Soc. 1945;37: 81–91.

16. Amari S, Nagaoka H. Methods of information geometry. Providence (RI): American Mathematical Society and Oxford University Press; 2000.

17. Olsen SR, Bhandawat V, Wilson RI. Divisive normalization in olfactory population codes. Neuron. 2010;66: 287–299.

18. Bhandawat V, Olsen SR, Gouwens NW, Schlief ML, Wilson RI. Sensory processing in the *Drosophila* antennal lobe increases reliability and separability of ensemble odor representations. Nat Neurosci. 2007;10: 1474–1482.

19. Stevens CF. What the fly’s nose tells the fly’s brain. Proc Natl Acad Sci U S A. 2015;112: 9460–9465.

20. Stevens CF. A statistical property of fly odor responses is conserved across odors. Proc Natl Acad Sci U S A. 2016;113: 6737–6742.

21. Zwicker D, Murugan A, Brenner MP. Receptor arrays optimized for natural odor statistics. Proc Natl Acad Sci U S A. 2016;113: 5570–5575.

22. Qin S, Li Q, Tang C, Tu Y. Optimal compressed sensing strategies for an array of nonlinear olfactory receptor neurons with and without spontaneous activity. Proc Natl Acad Sci U S A. 2019;116: 20286–20295.

23. Kurtovic A, Widmer A, Dickson BJ. A single class of olfactory neurons mediates behavioural responses to a *Drosophila* sex pheromone. Nature. 2007;446: 542–546.

24. van der Goes van Naters W, Carlson JR. Receptors and neurons for fly odors in *Drosophila*. Curr Biol. 2007;17: 606–612.

25. Litwin-Kumar A, Harris KD, Axel R, Sompolinsky H, Abbott LF. Optimal degrees of synaptic connectivity. Neuron. 2017;93: 1153–1164.

26. Panzeri S, Senatore R, Montemurro MA, Petersen RS. Correcting for the sampling bias problem in spike train information measures. J Neurophysiol. 2007;98: 1064–1072.

27. Kraskov A, Stögbauer H, Grassberger P. Estimating mutual information. Phys Rev E Stat Nonlin Soft Matter Phys. 2004;69: 066138.

28. Ross BC. Mutual information between discrete and continuous data sets. PLoS One. 2014;9: e87357.

29. Stensmyr MC, Dweck HK, Farhan A, Ibba I, Strutz A, Mukunda L, et al. A conserved dedicated olfactory circuit for detecting harmful microbes in *Drosophila*. Cell. 2012;151: 1345–1357.

30. Cover TM, Thomas JA. Elements of information theory. 2nd ed. Hoboken (NJ): Wiley; 2006.

31. Green DM, Swets JA. Signal detection theory and psychophysics. New York: Wiley; 1966.

32. Macmillan NA, Creelman CD. Detection theory: A user’s guide. 2nd ed. Mahwah (NJ): Lawrence Erlbaum Associates; 2005.

33. Wright GA, Smith BH. Different thresholds for detection and discrimination of odors in the honey bee (*Apis mellifera*). Chem Senses. 2004;29: 127–135.

34. Cao LH, Yang D, Wu W, Zeng X, Jing BY, Li MT, et al. Odor-evoked inhibition of olfactory sensory neurons drives olfactory perception in *Drosophila*. Nat Commun. 2017;8: 1357.

35. Su CY, Menuz K, Reisert J, Carlson JR. Non-synaptic inhibition between grouped neurons in an olfactory circuit. Nature. 2012;492: 66–71.

36. Mantel N. The detection of disease clustering and a generalized regression approach. Cancer Res. 1967;27: 209–220.

37. Lin CC, Potter CJ. Re-classification of *Drosophila melanogaster* trichoid and intermediate sensilla using fluorescence-guided single sensillum recording. PLoS One. 2015;10: e0139675.

38. Ebrahim SAM, Dweck HKM, Weiss BL, Carlson JR. A volatile sex attractant of tsetse flies. Science. 2023;379: eade1877.

39. Dweck HK, Ebrahim SA, Thoma M, Mohamed AA, Keesey IW, Trona F, et al. Pheromones mediating copulation and attraction in *Drosophila*. Proc Natl Acad Sci U S A. 2015;112: E2829–E2835.

40. DeWeese MR, Meister M. How to measure the information gained from one symbol. Network. 1999;10: 325–340.

41. Chahda JS, Soni N, Sun JS, Ebrahim SAM, Weiss BL, Carlson JR. The molecular and cellular basis of olfactory response to tsetse fly attractants. PLoS Genet. 2019;15: e1008005.

42. den Otter CJ, van der Goes van Naters WM. Single cell recordings from tsetse (*Glossina m. morsitans*) antennae reveal olfactory, mechano- and cold receptors. Physiol Entomol. 1992;17: 33–42.

43. Voskamp KE, van der Goes van Naters WM, den Otter CJ. Comparison of single cell sensitivities to attractants in the tsetse *Glossina fuscipes fuscipes*, *G. morsitans morsitans* and *G. pallidipes*. Med Vet Entomol. 1999;13: 460–462.

44. Averbeck BB, Latham PE, Pouget A. Neural correlations, population coding and computation. Nat Rev Neurosci. 2006;7: 358–366.

45. Costa SIR, Santos SA, Strapasson JE. Fisher information distance: A geometrical reading. Discrete Appl Math. 2015;197: 59–69.

46. Torgerson WS. Multidimensional scaling: I. Theory and method. Psychometrika. 1952;17: 401–419.

47. Paninski L. Estimation of entropy and mutual information. Neural Comput. 2003;15: 1191–1253.

48. Vinje WE, Gallant JL. Sparse coding and decorrelation in primary visual cortex during natural vision. Science. 2000;287: 1273–1276.

49. Couto A, Alenius M, Dickson BJ. Molecular, anatomical, and functional organization of the *Drosophila* olfactory system. Curr Biol. 2005;15: 1535–1547.

50. Ward JH Jr. Hierarchical grouping to optimize an objective function. J Am Stat Assoc. 1963;58: 236–244.

